# *BDNF Val66Met* moderates the relationship between cardiorespiratory fitness and memory in cognitively normal older adults

**DOI:** 10.1101/408955

**Authors:** Belinda M. Brown, Natalie Castalanelli, Stephanie R. Rainey-Smith, James Doecke, Michael Weinborn, Hamid R. Sohrabi, Simon M. Laws, Ralph N Martins, Jeremiah J Peiffer

## Abstract

Higher cardiorespiratory fitness has been associated with enhanced cognitive function in older adults; yet, this relationship demonstrates a degree of variability. Thus, it is hypothesised that variation in genetic factors may influence the relationship between fitness and cognitive health. In this study we evaluate whether the *BDNF* Val66Met polymorphism moderates the relationship between cardiorespiratory fitness and verbal and visuospatial memory. Data from ninety-nine cognitively normal men and women aged 60 – 80 years were used. Fitness was assessed by peak oxygen consumption, and verbal and visuospatial memory were evaluated using well-validated measures. Participants were categorised into: lower-fit Met carriers, higher-fit Met carriers, lower-fit Val/Val, or higher-fit Val/Val. A significant interaction was observed between *BDNF* Val66Met and fitness on visuospatial memory performance; whereby lower-fit Met carriers performed 1SD lower than higher-fit Met carriers (*p*=0.04). We observed higher levels of fitness mitigated the deleterious effect of *BDNF* Met allele carriage on visuospatial memory. Future intervention studies should evaluate the effect of structured exercise on cognitive health between *BDNF* Val66Met carriers and Val/Val homozygotes.

## 1. Introduction

In older adults, higher levels of habitual physical activity have been consistently associated with enhanced cognitive performance and decreased cognitive decline in observational studies (Brown et al., 2012; Middleton et al., 2011; Schuit et al., 2001; Zhu et al., 2017); however, these findings are less consistent in randomised controlled trials (Young et al., 2015). This level of inconsistency highlights the complex interplay between physical activity and cognitive health, which genetic factors likely mediate. For instance, structured physical activity (i.e. exercise) upregulates the release of brain-derived neurotrophic factor (BDNF) which is involved in synaptic plasticity, neurogenesis and neuronal survival (Erickson et al., 2011). Within the brain, BDNF is at its highest levels in the hippocampus, a region vital for learning and memory function (Caccamo et al., 2010). Importantly, a single nucleotide polymorphism (SNP) of the *BDNF* gene involving a methionine (Met) substitution for valine (Val) at codon 66 (named the *BDNF* Val66Met; rs6265) can negatively impact activity-dependent secretion and intracellular trafficking of the BDNF protein, and is subsequently associated with reduced hippocampal neurogenesis (Egan et al., 2003). Carriers of the Met allele have been observed to have smaller hippocampal volume and poorer memory performance, than those that are homozygous for Val (wild-type; Kambeitz et al., 2012; Lim et al., 2014; Molendijk et al., 2012).

Animal (Nigam et al., 2017; Oliff et al., 1998) and human studies (Griffin et al., 2011; Vaynman et al., 2004) have consistently demonstrated an induction of BDNF following exercise, with a dose-dependent effect of intensity observed (Dalise et al., 2017). Not surprisingly, given the beneficial actions of BDNF on the hippocampus, increases in memory following exercise interventions have been demonstrated (Leckie et al., 2014). It is likely that *BDNF* Val66Met plays a moderating role in this relationship; yet, previous literature in this field is inconsistent. Erickson and colleagues (2013) described higher levels of self-reported physical activity as mitigating the negative effects of carrying a Met allele on working memory, but not other measures of cognition, such as episodic memory or visuospatial memory. Conversely, Canivet et al (2015) observed better episodic memory in physically active Val/Val homozygotes, compared with inactive Val/Val homozygotes, with no differences in episodic memory observed in active Met carriers versus inactive Met carriers. It is possible the above described studies reflect a moderating effect of *BDNF* Val66Met, as higher levels of physical activity may strengthen the beneficial impact of Val/Val homozygosity, and carriers of the Met allele may have suboptimal memory performance mitigated by physical activity.

Within the current literature, the use of self-reported physical activity may partially explain the conflicting evidence for the moderating effect of *BDNF* Val66Met. Indeed, self-reported measures of physical activity may be prone to over-, or under-, reporting in older adults (Dyrstad et al., 2014). Objective measures, such as activity trackers, can provide more accurate data (Downs et al., 2014; Dyrstad et al., 2014); however, a level of ecological inaccuracy may exist as individuals may increase their physical activity levels during the relatively short period of data collection (i.e. one to two weeks). Cardiorespiratory fitness (VO_2peak_) is associated with the amount and intensity of physical activity (Joyner and Lundby, 2018) an individual undertakes over a prolonged period of time (DeFina et al., 2015). Therefore, the use of VO_2peak_ in determining the influence of physical activity on cognition may provide a more stable marker for such an analyses, and should be considered, when appropriate.

In the current study we evaluated the moderating effect of the *BDNF* Val66Met polymorphism on the relationship between cardiorespiratory fitness and performance on hippocampal-dependent cognitive tasks in a cohort of cognitively normal older adults. Based on previous literature we formulated two parallel hypotheses: 1. Higher fitness (fit) will attenuate poorer memory function in Met carriers; and 2. Higher fitness will amplify positive effects on memory in Val/Val homozygotes. Similarly, we hypothesised that we will observe mean memory performance in the following order: Lower-fit Met carriers < Lower-fit Val/Val homozygotes < Higher-fit Met carriers < Higher-fit Val/Val homozygotes.

## 2. Methods

### 2.1 Participants

Data from ninety-nine community dwelling cognitively normal men and women aged 60 – 80 years old from the baseline assessment of the Intense Physical Activity and Cognition (IPAC) study were utilised for the current study. Details of the IPAC study, including recruitment, power analysis, inclusion and exclusion criteria, procedures and ethical considerations have been detailed previously (Brown et al., 2017). Briefly, all participants were cognitively normal and did not have any uncontrolled medical conditions known to influence cognitive function. All procedures were approved by the human research ethics committees of Edith Cowan University and Murdoch University, Western Australia. All participants provided written informed consent prior to data collection.

### 2.2 Cardiorespiratory fitness testing

Participants were assessed for their peak aerobic capacity (VO_2peak_) using a cycling based graded exercise test completed on an electromagnetically braked cycle ergometer (Velotron; RacerMate, USA). All tests used two-minute stage durations with consistent increases in work rate at each stage until participants reached volitional fatigue. To ensure similar test durations and stage progression, test selection was determined relative to the participants baseline body mass using the following criteria; 1) participants under 70 kg commence testing at 30 W with increases of 20 W at each stage, 2) participants between 70 and 100 kg commence testing at 30 W with increases of 25 W at each stage and 3) participants over 100 kg commence testing at 40 W with increases of 35 W at each stage. During each test, heart rate was continuously recorded and expired ventilation was collected and analysed as 15 second mean values, using a Parvo TrueOne (ParvoMedics, USA) metabolic cart, for the rate of oxygen consumption (VO_2_) and carbon dioxide production (VCO_2_). At test completion, maximal heart rate was determined as the highest value recorded during the test, and VO_2peak_ was determined as the highest 15 second mean VO_2_ value obtained during the final 2 minutes of the test. Additional criteria for the assessment of VO_2peak_ involve participants reaching a maximal heart rate greater than 85% of their age predicted maximum (i.e. (220 – age) * 0.85) and a respiratory exchange ratio (VCO_2_/VO_2_) greater than 1.15.

Using the VO_2peak_ data, cut-off points were used to categorise our cohort into *higher-fit* and *lower-fit* groups. This terminology does not reflect cut-offs that are based on improved cognitive health outcomes (as this remains unknown), rather we employed age and gender based cut-offs from the 50^th^ percentile of the *Fitness Registry and the Importance of Exercise: A National Data Base* (FRIEND; Kaminsky et al., 2015). We utilised the median from the larger FRIEND cohort (*n* = 1236, of relevant age), rather than our own, as our groupings were small (e.g. *n =* 22 for men aged 70-79), and thus the median of our grouping may not reflect that of the wider community. The VO2peak cut-off points utilised within the current study are as follows (above these cut-offs categorised as *higher-fit*, equal to or below these cut-offs categorised as *lower-fit*): men aged 60-69y: 28.2 mL·kg^-1^·min^-1^; men aged 70-79y: 24.4 mL·kg^-1^·min^-1^; women aged 60-69y: 20.0 mL·kg^-1^·min^-1^; women aged 70-79y: 18.3 mL·kg^-1^·min^-1^.

### 2.3 Genotyping

Genotyping was completed as per manufacturer’s instructions. Briefly, DNA was extracted from whole blood aliquots using QIAamp DNA Blood Maxi Kits (Qiagen, Hilden, Germany). TaqMan genotyping assays were used to determine *APOE* genotype (rs7412, assay ID: C____904973_10; rs429358, assay ID: C____3084793_20), and *BDNF* Val66Met single nucleotide polymorphism (rs6265, assay ID: C____11592758_10). Within our cohort we observed Val/Val homozygotes, *n* = 61; Val/Met heterozygotes, *n* = 28; and Met/Met homozygotes, *n* = 3. This genotypic distribution did not deviate from Hardy-Weinberg Equilibrium (*p* = 0.45). Due to the low frequency of Met/Met homozygotes, a single group of Met carriers (*n* = 31) was created; consistent with previous studies (Erickson et al., 2013; Lim et al., 2013).

### 2.4 Cognitive testing

Hippocampal-dependent tasks were identified within the larger IPAC study cognitive battery for investigation within this project. More specifically, we examined auditory verbal memory using the California Verbal Learning Test-2^nd^ edition (CVLT-2) long delay recall (Delis, 2000), and visuospatial memory using the Brief Visual Memory Test (BVMT) long delay recall (Benedict, 1997) and the Continuous Paired Associate Learning Task (CPAL) errors from Cogstate (Maruff et al., 2009).

### 2.5 Statistical analysis

All statistical analysis was undertaken using SPSS version 24 (IBM Corporation, Armonk, NY). All cognitive data was within the acceptable range of skewness and kurtosis (within -2 and +2). Statistical significance was set at *p* < 0.05. Missing data for cardiorespiratory fitness (*n* = 3) was assessed and found to be missing at random (Little MCAR, *p* > 0.10). Subsequently, this data was imputed by multiple imputations using predictive mean matching (20 imputations and 200 iterations).

Independent samples t-tests and chi-square analyses were used to evaluate differences in continuous and categorical variables, respectively, across the fitness and *BDNF* Val66Met groups. Analysis of covariance (ANCOVA) was conducted with cognitive test scores entered as the dependent variables (into separate models) and *BDNF* Val66Met polymorphism (Val/Val homozygote or Met carrier), fitness group (lower-fit or higher-fit), age, gender, years of education and body mass index (BMI), and a *BDNF* Val66Met*fitness group interaction entered into the models. In order to test our hypotheses, parameter estimates were examined in the models that returned a significant *BDNF* Val66Met*fitness group interaction: A four group variable (lower-fit Met carriers, lower-fit Val/Val homozygotes, higher-fit Met carriers and higher-fit Val/Val homozygotes) was created and subsequently entered into an additional ANCOVA in place of the *BDNF* polymorphism, fitness, and the *BDNF* Val66Met*fitness interaction (all other variables remained).

## 3. Results

The higher-fit group was significantly younger (67.0 ± 4.3 y) than the lower-fit group (71.5 ± 4.4 y; *t* = 4.67, *p* < 0.001; Table 1). Furthermore, the higher-fit group had a lower BMI (24.4 ± 3.0 kg.m^2^), than the lower-fit group (27.1 ± 3.8 kg.m^2^; *t* = 3.88, *p* < 0.001). No differences were observed between the *BDNF* Met carriers and Val/Val homozygotes in terms of the descriptive and demographic variables.

**Table 1:** Descriptive statistics of the total cohort, and breakdown by fitness group and BDNF Val66Met carriage. Data presented as mean ± standard deviation, unless otherwise noted.

|  | Total<br>(*n* = 99) | Cardiorespiratory fitness |  |  | BDNF Val66Met polymorphism |  |  |
| --- | --- | --- | --- | --- | --- | --- | --- |
|  |  | Lower-fit<br>(*n* = 46) | Higher-fit<br>(*n* = 53) | p-value | Val/Val<br>(*n* = 61) | Met carriers<br>(*n* = 38) | p-value |
| Age, years | 69.1 ± 5.2 | 71.5 ± 5.1 | 67.0 ± 4.3 | <0.001 | 69.3 ± 5.3 | 68.8 ± 5.1 | 0.61 |
| Gender, % (*n*) female | 54.5 (54) | 54.3 (25) | 54.7 (29) | 0.97 | 50.8 (31) | 60.5 (23) | 0.35 |
| Education, years | 14.0 ± 2.3 | 13.7 ± 2.4 | 14.3 ± 2.1 | 0.18 | 13.9 ± 2.3 | 14.3 ± 2.3 | 0.41 |
| *APOE* ε4 allele carriage, % (*n*) | 26.3 (26) | 17.4 (8) | 34.0 (18) | 0.06 | 27.9 (17) | 23.7 (9) | 0.64 |
| *BDNF* Val66Met carriage, % (*n*) | 38.3 (38) | 39.1 (18) | 37.7 (20) | 0.89 |  |  |  |
| VO<sub>2peak</sub>, mL·kg<sup>-1</sup>·min<sup>-1</sup> | 23.0 ± 6.5 | 18.2 ± 4.2 | 27.2 ± 5.2 | <0.001 | 23.4 ± 7 | 22.4 ± 5.7 | 0.49 |
| BMI, kg/m<sup>2</sup> | 25.7 ± 3.7 | 27.1 ± 3.8 | 24.4 ± 3.0 | <0.001 | 25.6 ± 3.9 | 25.8 ± 3.2 | 0.75 |

Differences in groups determined by chi-square analysis (categorical variables) and independent sample t-tests (continuous variables).
Abbreviations: *APOE* ε4, apolipoprotein E epsilon 4; *BDNF* Val66Met, brain-derived neurotrophic factor Valine66Methionine single nucleotide polymorphism; BMI, body mass index; fit, fitness; VO<sub>2peak</sub>, peak aerobic capacity (fitness measurement).

No differences were observed between the *BDNF* Met carriers and Val/Val homozygotes in terms of performance on the CVLT delayed recall (*F* = 0.07, *p* = 0.79), BVMT delayed recall (*F* = 0.17, *p* = 0.68) and CPAL (*F* = 0.84, *p* = 0.36; Table 2). No differences between the lower-fit and higher-fit groups were observed on the CVLT delayed recall (*F* = 0.73, *p* = 0.40) and BVMT delayed recall (*F* = 0.05, *p* = 0.83); however, the higher-fit group performed better than the lower-fit group (in terms of less errors) on the CPAL task (*F* = 7.94, *p* = 0.006). A significant effect of the *BDNF* Val66Met*fitness interaction was observed on CPAL performance (*F* = 4.33, *p* = 0.04). Following the creation of the four BDNF/fitness groups, the ANCOVAs were re-run for the CPAL, and parameter estimates examined (mean differences between groups shown in Figure 1): the lower-fit Met carrier group performed worse than the higher-fit Met carriers (*B* = -60.9, *p* = 0.001), lower-fit Val/Val homozygotes (*B* = -34.7, *p* = 0.04), and higher-fit Val/Val homozygotes (*B* = -47.3, *p* = 0.007) groups.

**Figure 1.**
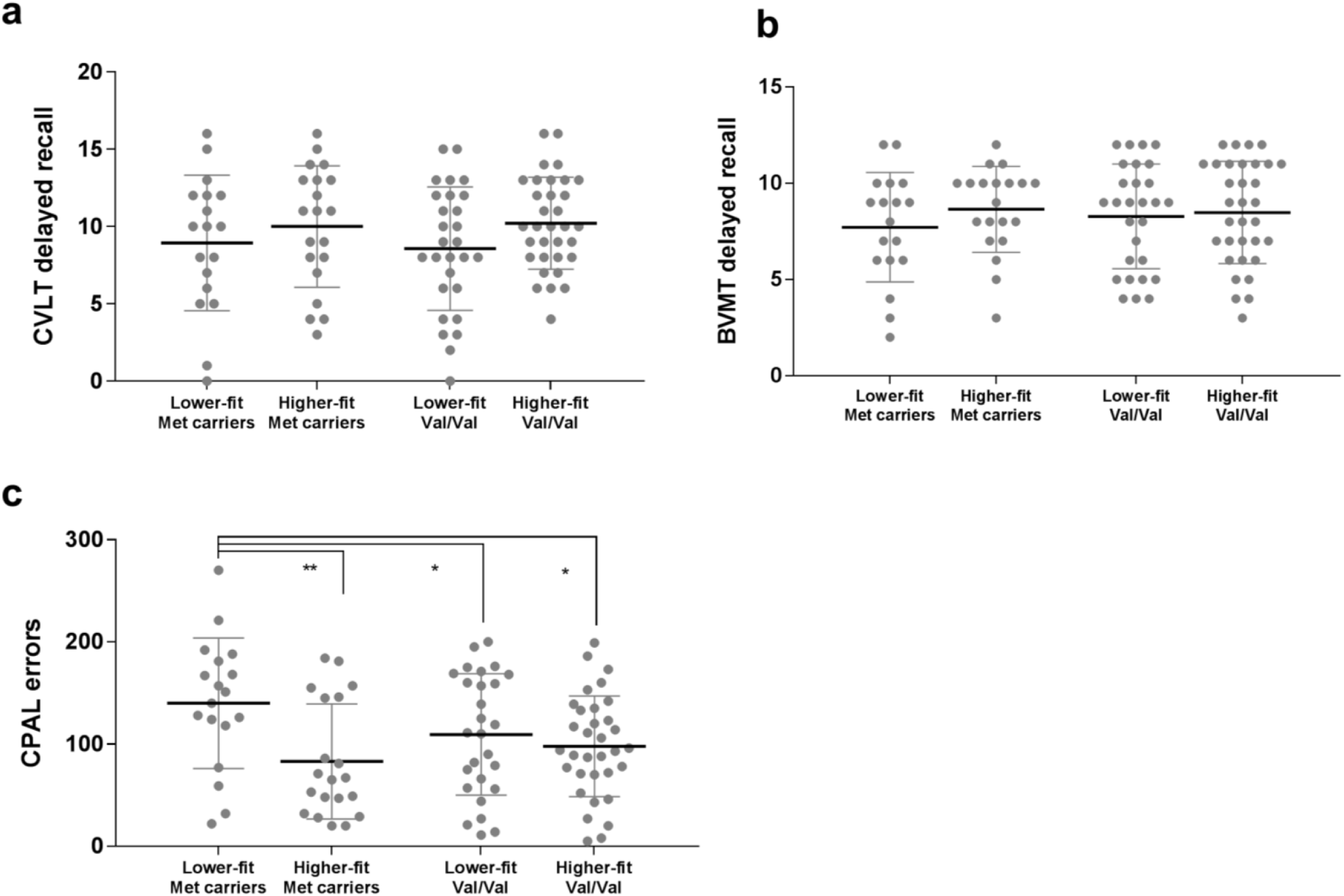
The relationship between the fitness and BDNF Val66Met interaction on memory performance. Lower-fit Met carriers scored significantly more errors on the CPAL task (c), compared with all other groups. No differences were observed across the groups on the CVLT delayed recall (a) and BVMT delayed recall (b). Raw means and standard deviations shown, with p-value indicators from analysis of covariance, with lower-fit Met carriers set as the reference group. * p < 0.05, **p < 0.01. Abbreviations: BVMT, Brief Visual Memory Test; CPAL, Continuous Paired Associates Learning Task, CVLT, California Verbal Learning Test; fit, fitness; Met, methionine; val, valine.

**Table 2:** Analysis of covariance examining differences in verbal and visuospatial memory performance between individuals classed as lower-fit versus higher-fit and BDNF Val66Met carriers and Val/Val homozygotes.

|  | Age | Gender | Education | BMI | Fitness |  |  | BDNF Val66Met |  |  | BDNF Val66Met x fitness |
| --- | --- | --- | --- | --- | --- | --- | --- | --- | --- | --- | --- |
| | *F* | *F* | *F* | *F* | Lower-fit<br>Mean ± SE<sup>†</sup> | Higher-fit<br>Mean ± SE<sup>†</sup> | *F* | Val/Val<br>Mean ± SE<sup>†</sup> | Met carriers<br>Mean ± SE<sup>†</sup> | *F* | *F* |
| CVLT delayed recall | 1.81 | 10.92** | 0.67 | 0.81 | 8.98 ± 0.59 | 9.69 ± 0.53 | 0.73 | 9.44 ± 0.46 | 9.22 ± 0.58 | 0.07 | 0.01 |
| BVMT delayed recall | 3.49 | 0.01 | 0.05 | 0.05 | 8.24 ± 0.44 | 8.38 ± 0.40 | 0.05 | 8.42 ± 0.34 | 8.19 ± 0.44 | 0.17 | 1.10 |
| CPAL task errors | 3.65 | 1.67 | 0.18 | 6.23* | 127.2 ± 9.10 | 90.4 ± 8.21 | 7.94** | 103.6 ± 7.1 | 114.1 ± 8.9 | 0.84 | 4.33* |

\* $p < 0.05$ , \*\* $p < 0.01$ . <sup>†</sup>Adjusted marginal means ± standard error. Abbreviations: BDNF Val66Met, brain-derived neurotrophic factor Valine66Methionine single nucleotide polymorphism; BMI, body mass index; BVMT, Brief Visual Memory Test; CPAL, Continuous Paired Associate Learning Task; CVLT, California Verbal Learning Test; fit, fitness; SE, standard error.

## 4. Discussion

In the current study we sought to examine the moderating effect of the *BDNF* Val66Met polymorphism on the relationship between cardiorespiratory fitness and performance on memory tasks. We proposed two hypotheses that would underlie the interaction between *BDNF* Val66Met and cardiorespiratory fitness on memory performance: Higher fitness levels would, 1) mitigate the deleterious effects of Met carriage on memory performance and, 2) be associated with enhanced memory performance beyond the beneficial effects of Val/Val homozygosity. In partial support of our hypothesis, we observed a significant effect of a *BDNF* Val66Met*fitness interaction on visuospatial memory. More specifically, we report higher-fit Met carriers performed significantly better than lower-fit Met carriers on the CPAL task, indicating a potential protective effect of higher cardiorespiratory fitness against the deleterious effects of Met carriage.

The 1SD difference on the CPAL task observed between the lower-fit and higher-fit Met carriers supports our proposed hypotheses that higher levels of cardiorespiratory fitness can contribute to modifying the negative effects of *BDNF* Met carriage. These findings support those reported by Erickson et al (2013), who observed greater amounts of self-reported habitual physical activity attenuated poor working memory performance in Met carriers; however, these authors did not report an effect on visuospatial memory. The underlying mechanism of an effect of fitness on memory in Met carriers remains to be elucidated. It is possible that higher levels of fitness are associated with greater *BDNF* gene expression, in Met carriers only. Alternatively, Val/Val homozygotes may perform at a ‘ceiling’ level on memory tasks; and thus memory changes in response to increased fitness may be minimal. However, Canivet and colleagues (2015) observed no difference in memory in reportedly (i.e. self-reported) active compared with inactive Met carriers. It is probable that a number of factors are associated with the level of moderation provided by the *BDNF* Val66Met polymorphism on the relationship between fitness/physical activity and memory. Such factors likely include the cognitive tasks utilised, age of the cohort, sample size, level of cognitive functioning of the cohort and the measure of fitness/exercise. While physical activity is associated with aerobic fitness (Bastone Ade et al., 2015; Joyner and Lundby, 2018; Stofan et al., 1998), heterogeneity in the response of aerobic fitness to physical activity has been observed (Bouchard et al., 1999). It is therefore possible that categorising cohorts based on aerobic fitness alone could represent much larger differences in physical activity levels (Joyner and Lundby, 2018); thus, explaining the difference in outcomes observed in our study compared with Canivet et al (2015).

Our data does not support our second hypothesis that greater fitness would be associated with enhanced cognitive performance beyond the influence of Val/Val homozygosity. Indeed, no differences were noted in any of the memory measures between the high and low fitness groups in the Val/Val cohort (Table 2). This finding is again consistent with those of Erikson et al. (2013) and in contrast to Canivet et al. (2015). Should our measure of aerobic fitness accurately represent physical activity, as outlined above, then our findings support that high levels of physical activity do not enhance cognitive performance in Val/Val homozygotes, at least for the cognitive domains that were assessed. Of note however, our study, as well as those of Erikson et al and Canivet et al, utilised cognitively normal cohorts. It is possible that in such cohorts, the influence of *aerobic fitness* is only observed in the presence of the detrimental Met allele. From the present data, we cannot confirm or refute this hypothesis. As such, a study examining the impact of aerobic fitness on cognitive performance between Met carriers and Val/Val homozygotes in both cognitively normal individuals and those with cognitive decline, is needed.

Our data are supported by a large body of animal work suggesting that improvements in memory following exercise are mediated by BDNF (Vaynman et al., 2004). For instance, animal running is associated with increased expression levels of BDNF mRNA, and by blocking the binding of BDNF to its receptor, TrkB, the beneficial effects to cognitive health dissipate. In *BDNF* Met carriers the prodomain structure of BDNF is altered, which results in improper protein folding and intracellular trafficking of mature BDNF. Thus, in Met carriers, there is an increased binding of the precursor protein to its receptor, p75, and reduced binding of mature BDNF to TrkB. Binding of precursor BDNF to p75 is associated with an induction of apoptosis (Teng et al., 2005), whereas binding of mature BDNF to TrkB induces the beneficial effects associated with BDNF such an enhanced neurogenesis and neuronal survival. To-date, it is unclear as to how physical activity could modulate BDNF action in Met carriers. Physical activity could act upon protein folding in Met carriers, leading to increased levels of mature BDNF, or alternatively it could reduce apoptosis associated with precursor BDNF-p75 binding. Nevertheless, it is important to note that our study did not utilise any measures of BDNF protein levels, and thus our conclusions regarding the role of the BDNF protein as a mediator in the relationship between fitness and memory are merely speculative.

The appropriateness of the cognitive measures utilised within the current study also should be considered, as we only observed an effect of the fitness\**BDNF* Val66Met interaction on the CPAL (visuospatial memory), but not the BVMT (also visuospatial memory) nor the CVLT (a measure of verbal memory). There is great inconsistency across previous studies examining the relationship between exercise and various cognitive domains, with generally small effect sizes reported (Kelly et al., 2014). Within the current study, we selected three hippocampal-dependent cognitive tasks; yet, it could be hypothesised that an effect of the fitness\**BDNF* interaction was observed for only one visuospatial memory task, given its greater dependency on hippocampal function. The CPAL task comprises two highly hippocampal-dependent cognitive factors, visuospatial function and memory; it is possible that tasks that are *most* dependent on the hippocampus (i.e. spatial tasks over verbal tasks), receive the greatest moderating effect of the *BDNF* polymorphism. It is however also important to note that ∼11% of the cohort performed at ceiling (score of 12) on the BVMT long delay recall, whilst no participants performed at ceiling on the CPAL (score of 0 errors). Thus, it is possible that the CPAL is more effective at detecting subtle differences in visuospatial memory performance in cognitively normal individuals; however, this would need to be validated in a large longitudinal study.

The examination of *BDNF* Val66Met as a moderating genetic factor in the relationship between fitness and memory is based on robust biological evidence of the effect of exercise on brain health. Nevertheless, it is likely that *BDNF* Val66Met is not the only genetic factor contributing to the relationship. For instance, the *APOE* ε4 allele is known to moderate the relationship between physical activity, and levels of beta-amyloid (the neurotoxic protein implicated in Alzheimer’s disease; Brown et al., 2013; Head et al., 2012). Higher levels of self-reported physical activity have been shown to reduce the negative effect of carrying an *APOE* ε4 allele on brain Aβ-amyloid burden. The *BDNF* Val66Met polymorphism and *APOE* likely represent different underlying mechanisms in the relationship between exercise and cognitive health. As such, future research is needed to explore genetic moderators of brain health to establish a comprehensive knowledge base that could be utilised for the development of individualised exercise regimens based on genetic factors.

The results reported here should be interpreted within the context of the limitations of the current study. Importantly, the current study was cross-sectional, and a causal direction cannot be interpreted. It is also possible that Met carriage and subsequently poorer memory is associated with lower levels of habitual physical activity, contributing to lower fitness levels. However, the mean performance on cognitive tasks by our Met carriers does not suggest function that is below a clinical threshold; thus, we deem it unlikely that physical activity, and subsequently cardiorespiratory fitness, levels would reduce before clinically significant cognitive decline is detected. Furthermore, our sample size, although sufficiently powered for the statistical analysis, was relatively small, particularly when grouping individuals by fitness and *BDNF* Val66Met variables. However, the use of an objective measure such as cardiorespiratory fitness, and in addition to the relatively low variability in our cognitive task results, we are confident in our findings.

## Conclusions

Our results support previous literature to suggest an effect of a *BDNF* Val66Met*fitness interaction on visuospatial memory. Our data supports the notion that increased cardiorespiratory fitness is protective against the deleterious effects of carrying the Met allele. Nevertheless, this association requires further investigation, ideally within large randomised controlled exercise trials, where the benefits of exercise interventions on cognitive performance can be compared between those who are *BDNF* Val66Met carriers and those who are Val/Val homozygotes.

## Acknowledgements

All authors would like to thank the IPAC study participants and their families for their generous participation.

---

**Supplementary Table 1.** Descriptive statistics of fitness and BDNF Val66Met carriage groupings. Data presented as mean ± standard deviation, unless otherwise noted.

|  | Lower-fit Met carriers<br>(*n* = 18) | Higher-fit Met carriers<br>(*n* = 20) | Lower-fit Val/Val<br>(*n* = 28) | Higher-fit Val/Val<br>(*n* = 33) |
| --- | --- | --- | --- | --- |
| Age, years | 69.8 ± 5.6 | 67.9 ± 4.5 | 72.6 ± 4.5<sup>a</sup> | 66.5 ± 4.2 |
| Gender, % (*n*) female | 61 (11) | 60 (12) | 50 (14) | 51 (17) |
| Education, years | 14 ± 2 | 15 ± 2 | 14 ± 2 | 14 ± 2 |
| *APOE* ε4 allele carriage %<br>(*n*) | 12.5 (2) | 35 (7) | 21.4 (6) | 33.3 (11) |
| VO<sub>2</sub>peak, mL·kg<sup>-1</sup>·min<sup>-1</sup> | 17.9 ± 3.8 | 26.5 ± 5.9 | 18.4 ± 4.4 | 27.6 ± 5.9 |
| BMI, kg/m<sup>2</sup> | 26.8 ± 3.4 | 24.9 ± 2.9 | 27.3 ± 4.1<sup>b</sup> | 24.2 ± 3.1 |

<sup>a</sup>Lower-fit Val/Val homozygotes were significantly older than both higher-fit Met carriers and higher-fit Val/Val homozygotes. <sup>b</sup>Lower-fit Val/Val homozygotes had significantly higher BMI than higher-fit Val/Val homozygotes. Differences in groups determined by chi-square analysis (categorical variables) and analysis of variance (continuous variables; post-hoc Tukey's). Abbreviations: *APOE* ε4, apolipoprotein E epsilon 4; BDNF Val66Met, brain-derived neurotrophic factor Valine66Methionine single nucleotide polymorphism; BMI, body mass index; fit, fitness; VO<sub>2</sub>peak, peak aerobic capacity (fitness measurement).

